# Genomic analysis reveals local transmission of SARS-CoV-2 in early pandemic phase in Peru

**DOI:** 10.1101/2020.09.05.284604

**Authors:** Carlos Padilla-Rojas, Karolyn Vega-Chozo, Marco Galarza-Perez, Henri Bailon Calderon, Priscila Lope-Pari, Johanna Balbuena-Torres, David Garcia Neyra, Maribel Huaringa-Nuñez, Nancy Rojas-Serrano, Omar A. Caceres-Rey

## Abstract

The dissemination of cases of the new SAR-COV-2 coronavirus represents a serious public health problem for Latin America and Peru. For this reason, it is important to characterize the genome of the isolates that circulate in Latin America. To characterize the complete genome of first samples of the virus circulating in Peru, we amplified seven overlapping segments of the viral genome by RT-PCR and sequenced using Miseq platform. The results indicate that the genomes of the Peruvian SARS-COV-2 samples belong to the genetic groups G and S. Likewise, a phylogenetic and MST analysis of the isolates confirm the introduction of multiple isolates from Europe and Asia that, after border closing, were transmitted locally in the capital and same regions of the country. These Peruvian samples (56%) grouped into two clusters inside G clade and share B.1.1.1 lineage. The characterization of these isolates must be considered for the use and design of diagnostic tools, and effective treatment and vaccine formulations.

## INTRODUCTION

A new coronavirus called SARS-COV-2 causes Severe Acute Respiratory Syndrome (SARS) that currently produced a pandemic. SARS-CoV-2 is a zoonotic virus of the beta coronavirus genus that belongs to SARS-type viruses (bat-SL-CoVZC45 and bat-SL-CoVZXC21) (1). These viruses usually cause mild respiratory infections in healthy adults and more severe and sometimes fatal effects in older adults and people with comorbidities (2). The SARS-CoV-2 virus is an enveloped virus with a 29.903 kb, positive sense, and single-stranded RNA genome. This genome is one of the largest among RNA viruses, and it encodes four structural proteins (E, M, N, and S), whose function is to protect the viral genome and 16 non-structural proteins (NSP) that regulate RNA synthesis and its processing, such as RNA dependent RNA polymerase (RdRp) (NSP12) and helicase (NSP13). Simultaneously, other proteins facilitate the function of viral enzymes, such as NSP7 and NSP10 (3).

In December 2019, the SARS-CoV-2 virus initially caused an outbreak of atypical pneumonia in Wuhan, China; since then, it has spread to almost all countries in the world. For this reason, the WHO declared it a pandemic on the 11th March, and later this new disease was named COVID-19 (4,5). WHO ranked the COVID-19 as the sixth international public health emergency of global concern after H1N1 (2009), Polio (2014), Ebola from West Africa (2014), Zika (2016), and Ebola from the Democratic Republic of the Congo (2019). Thus, due to the virus’s high transmission capacity, a growing incidence of infection, and the ability to be transmitted by asymptomatic carriers, and call on the world’s health systems to cooperate to prevent the spread of SARS-CoV-2 (6).s

At the date of this publication, the COVID-19 has caused more than 16 and a half million confirmed cases and more than 650,000 deaths worldwide. America is one of the most affected regions by the disease, with more than 8,800,000 affected patients, where the USA and Brazil are the most affected countries (4,263,531 and 2,442,375 confirmed cases respectively). Peru ranks seventh among the countries with the most confirmed cases worldwide (more than 600,000 infected), and first place in mortality rate (more than 28 000 deaths) (8). The first Peruvian case of COVID-19 was notified on 5^th^ March 2020 in a 25-year-old man who arrived in the country after visiting Spain, France, and the Czech Republic. Subsequently, new cases began to be notified in other cities of Peru in such a way that the Peruvian government decreed the international closure of borders and mandatory quarantine at the national level on 16^th^ March.

To control the COVID-19 burden, it is necessary to understand the genomic structure of the SARS-CoV-2 virus, identify new mutations associated with its transmission, and infer these transmission routes. Knowledge of these aspects will allow the development of vaccines, implement better disease control, and generate new diagnostic methods. In this context, our study seeks to contribute with genomic analysis of the early transmission of SARS-CoV-2 virus circulating in the beginning of the COVID-19 pandemics in Peru.

## RESULTS

We produced seven overlapping fragments of a size between 5 Kb to 6 Kb from the genome of SARS-COV-2 by RT-PCR (Figure 1). In some cases, we also get nonspecific amplification, eliminated during the fragment’s purification. We submitted 34 SARS-COV-2 genome sequences from Peru to the international Global Initiative on Sharing All Influenza Data (GISAID) database. We identified six SARS-CoV-2 genetic lineages (A2, A5, B.1, B1.1, B1.1.1, B1.5) according to GISAID and Pangolin analysis (Table 2). Most Peruvian SARS-CoV-2 sequences here obtained were classified as clade B.1 (38%, n = 13), and particularly within the sub-clade B.1.1.1 (29.4%, n = 10) (Table 2).

**Table 1.**
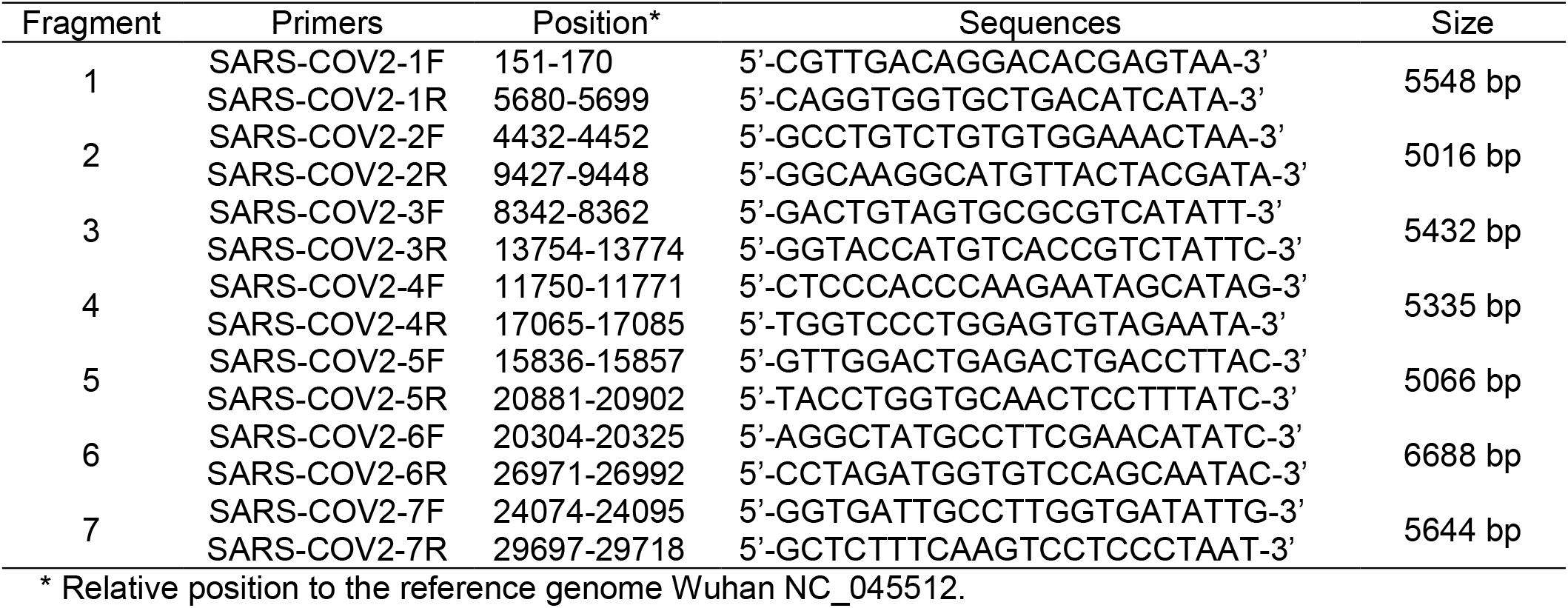
Primers designed for amplification of overlapping fragments of the SARS-COV-2 genome.

**Table 2.**
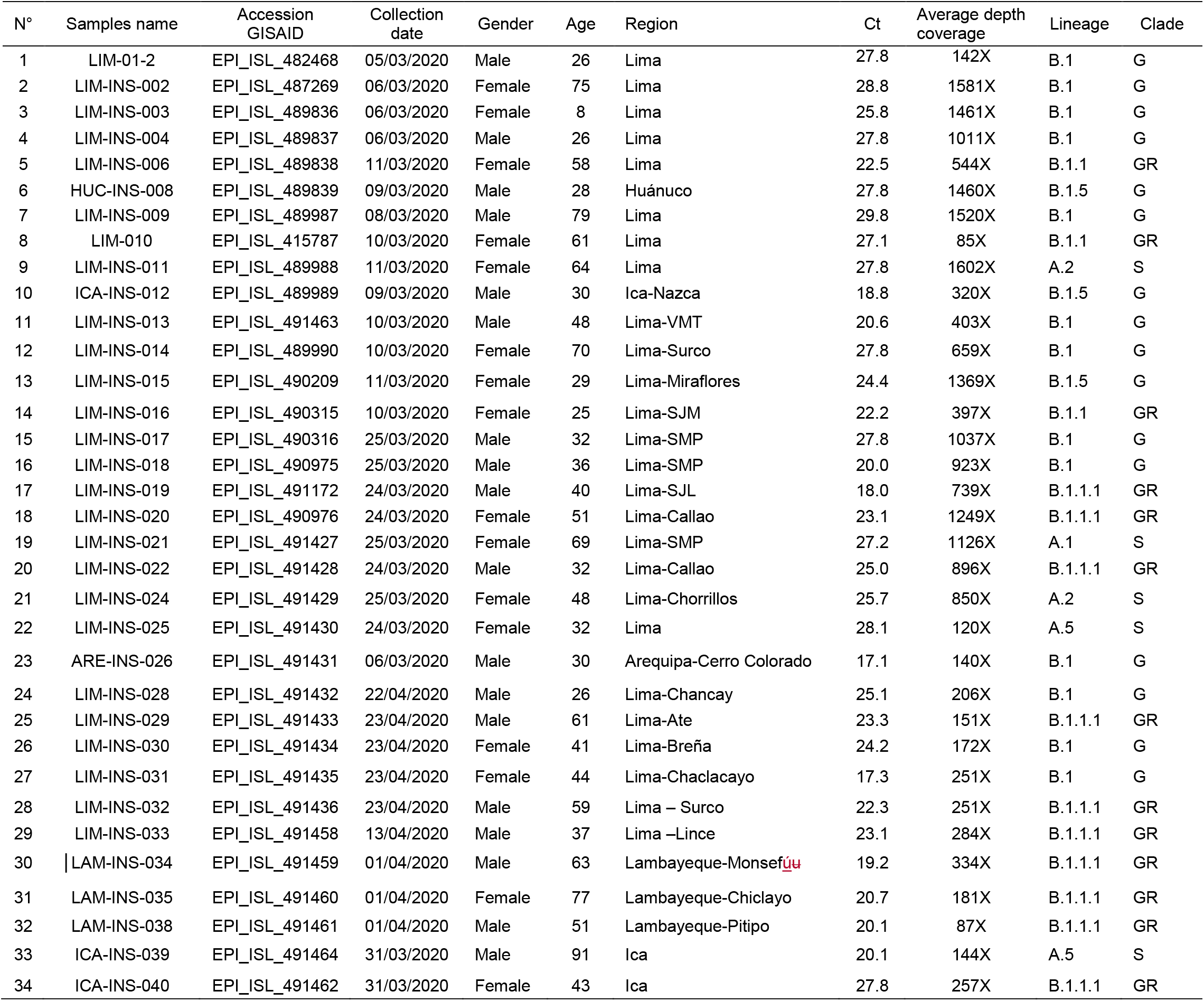
Samples analyzed by massive sequencing of circulating SARS-CoV-2 in Peru.

| N° | Samples name | Accession<br>GISAID | Collection<br>date | Gender | Age | Region | Ct | Average depth<br>coverage | Lineage | Clade |
| --- | --- | --- | --- | --- | --- | --- | --- | --- | --- | --- |
| 1 | LIM-01-2 | EPI_ISL_482468 | 05/03/2020 | Male | 26 | Lima | 27.8 | 142X | B.1 | G |
| 2 | LIM-INS-002 | EPI_ISL_487269 | 06/03/2020 | Female | 75 | Lima | 28.8 | 1581X | B.1 | G |
| 3 | LIM-INS-003 | EPI_ISL_489836 | 06/03/2020 | Female | 8 | Lima | 25.8 | 1461X | B.1 | G |
| 4 | LIM-INS-004 | EPI_ISL_489837 | 06/03/2020 | Male | 26 | Lima | 27.8 | 1011X | B.1 | G |
| 5 | LIM-INS-006 | EPI_ISL_489838 | 11/03/2020 | Female | 58 | Lima | 22.5 | 544X | B.1.1 | GR |
| 6 | HUC-INS-008 | EPI_ISL_489839 | 09/03/2020 | Male | 28 | Huánuco | 27.8 | 1460X | B.1.5 | G |
| 7 | LIM-INS-009 | EPI_ISL_489987 | 08/03/2020 | Male | 79 | Lima | 29.8 | 1520X | B.1 | G |
| 8 | LIM-010 | EPI_ISL_415787 | 10/03/2020 | Female | 61 | Lima | 27.1 | 85X | B.1.1 | GR |
| 9 | LIM-INS-011 | EPI_ISL_489988 | 11/03/2020 | Female | 64 | Lima | 27.8 | 1602X | A.2 | S |
| 10 | ICA-INS-012 | EPI_ISL_489989 | 09/03/2020 | Male | 30 | Ica-Nazca | 18.8 | 320X | B.1.5 | G |
| 11 | LIM-INS-013 | EPI_ISL_491463 | 10/03/2020 | Male | 48 | Lima-VMT | 20.6 | 403X | B.1 | G |
| 12 | LIM-INS-014 | EPI_ISL_489990 | 10/03/2020 | Female | 70 | Lima-Surco | 27.8 | 659X | B.1 | G |
| 13 | LIM-INS-015 | EPI_ISL_490209 | 11/03/2020 | Female | 29 | Lima-Miraflores | 24.4 | 1369X | B.1.5 | G |
| 14 | LIM-INS-016 | EPI_ISL_490315 | 10/03/2020 | Female | 25 | Lima-SJM | 22.2 | 397X | B.1.1 | GR |
| 15 | LIM-INS-017 | EPI_ISL_490316 | 25/03/2020 | Male | 32 | Lima-SMP | 27.8 | 1037X | B.1 | G |
| 16 | LIM-INS-018 | EPI_ISL_490975 | 25/03/2020 | Male | 36 | Lima-SMP | 20.0 | 923X | B.1 | G |
| 17 | LIM-INS-019 | EPI_ISL_491172 | 24/03/2020 | Male | 40 | Lima-SJL | 18.0 | 739X | B.1.1.1 | GR |
| 18 | LIM-INS-020 | EPI_ISL_490976 | 24/03/2020 | Female | 51 | Lima-Callao | 23.1 | 1249X | B.1.1.1 | GR |
| 19 | LIM-INS-021 | EPI_ISL_491427 | 25/03/2020 | Female | 69 | Lima-SMP | 27.2 | 1126X | A.1 | S |
| 20 | LIM-INS-022 | EPI_ISL_491428 | 24/03/2020 | Male | 32 | Lima-Callao | 25.0 | 896X | B.1.1.1 | GR |
| 21 | LIM-INS-024 | EPI_ISL_491429 | 25/03/2020 | Female | 48 | Lima-Chorrillos | 25.7 | 850X | A.2 | S |
| 22 | LIM-INS-025 | EPI_ISL_491430 | 24/03/2020 | Female | 32 | Lima | 28.1 | 120X | A.5 | S |
| 23 | ARE-INS-026 | EPI_ISL_491431 | 06/03/2020 | Male | 30 | Arequipa-Cerro Colorado | 17.1 | 140X | B.1 | G |
| 24 | LIM-INS-028 | EPI_ISL_491432 | 22/04/2020 | Male | 26 | Lima-Chancay | 25.1 | 206X | B.1 | G |
| 25 | LIM-INS-029 | EPI_ISL_491433 | 23/04/2020 | Male | 61 | Lima-Ate | 23.3 | 151X | B.1.1.1 | GR |
| 26 | LIM-INS-030 | EPI_ISL_491434 | 23/04/2020 | Female | 41 | Lima-Breña | 24.2 | 172X | B.1 | G |
| 27 | LIM-INS-031 | EPI_ISL_491435 | 23/04/2020 | Female | 44 | Lima-Chaclacayo | 17.3 | 251X | B.1 | G |
| 28 | LIM-INS-032 | EPI_ISL_491436 | 23/04/2020 | Male | 59 | Lima – Surco | 22.3 | 251X | B.1.1.1 | GR |
| 29 | LIM-INS-033 | EPI_ISL_491458 | 13/04/2020 | Male | 37 | Lima –Lince | 23.1 | 284X | B.1.1.1 | GR |
| 30 | LAM-INS-034 | EPI_ISL_491459 | 01/04/2020 | Male | 63 | Lambayeque-Monsefú | 19.2 | 334X | B.1.1.1 | GR |
| 31 | LAM-INS-035 | EPI_ISL_491460 | 01/04/2020 | Female | 77 | Lambayeque-Chiclayo | 20.7 | 181X | B.1.1.1 | GR |
| 32 | LAM-INS-038 | EPI_ISL_491461 | 01/04/2020 | Male | 51 | Lambayeque-Pitipo | 20.1 | 87X | B.1.1.1 | GR |
| 33 | ICA-INS-039 | EPI_ISL_491464 | 31/03/2020 | Male | 91 | Ica | 20.1 | 144X | A.5 | S |
| 34 | ICA-INS-040 | EPI_ISL_491462 | 31/03/2020 | Female | 43 | Ica | 27.8 | 257X | B.1.1.1 | GR |

**Figure 1.**
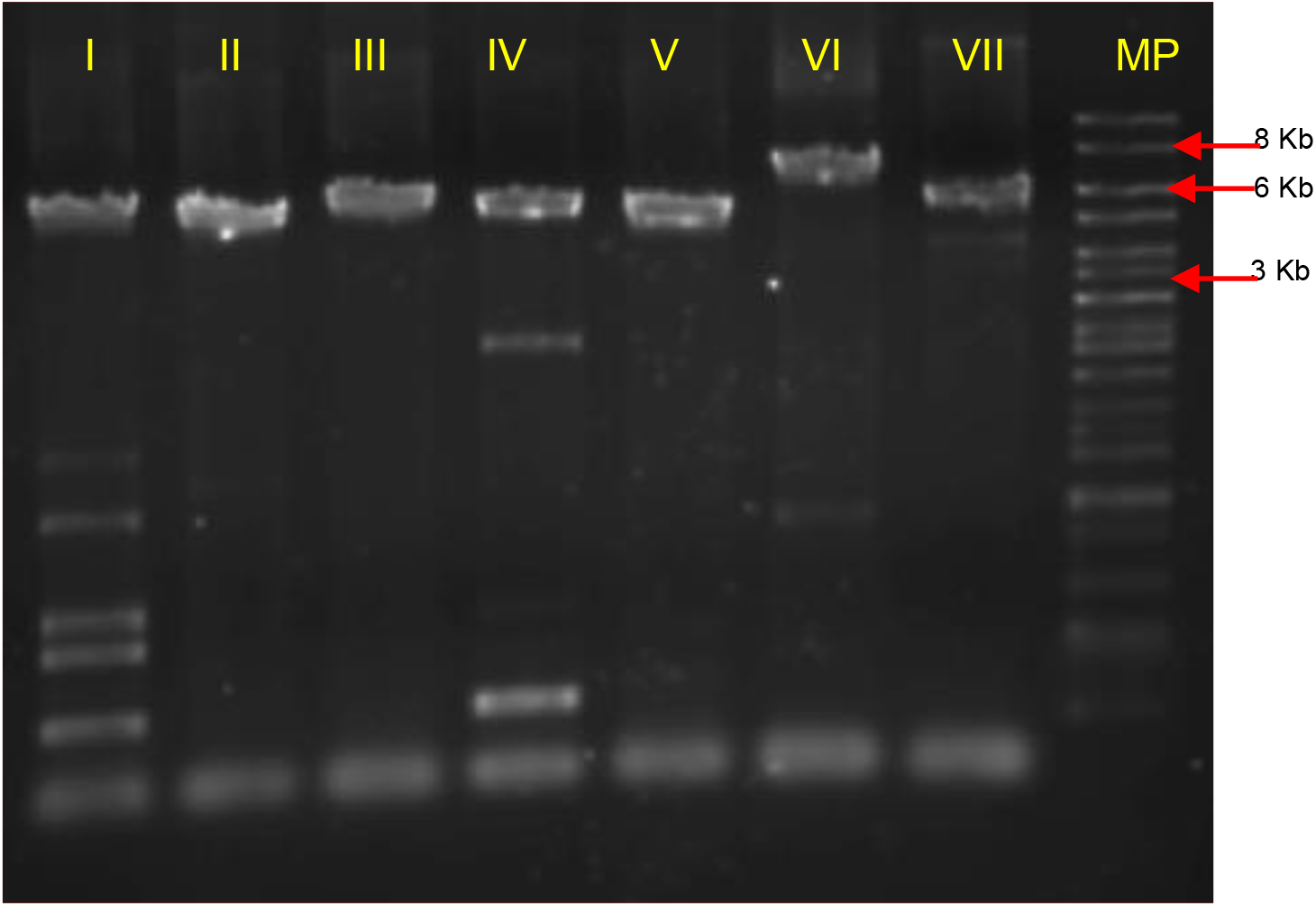
Amplified fragments of the SARS-COV-2 genome: I) 1, II) 2, III) 3, IV) 4, V) 5, VI) 6, and VII) 7. MP: molecular weight marker 1 Kb ladder (Thermo Scientific).

### Phylogenetic analysis

The genomes of the SARS-CoV-2 coronavirus from Peru, compared to the genomes of the viruses reported in different countries, were grouped into genetic clades G (85.3%) and S (14.7%), probably transmitted to people returning from Europe countries (such as Spain, Italy and Germany) and Asians countries (Taiwan). We also observed the presence of new Peruvian genotypes of the virus in early March and early April 2020, consistent with a local transmission (Figure 2).

**Figure 2.**
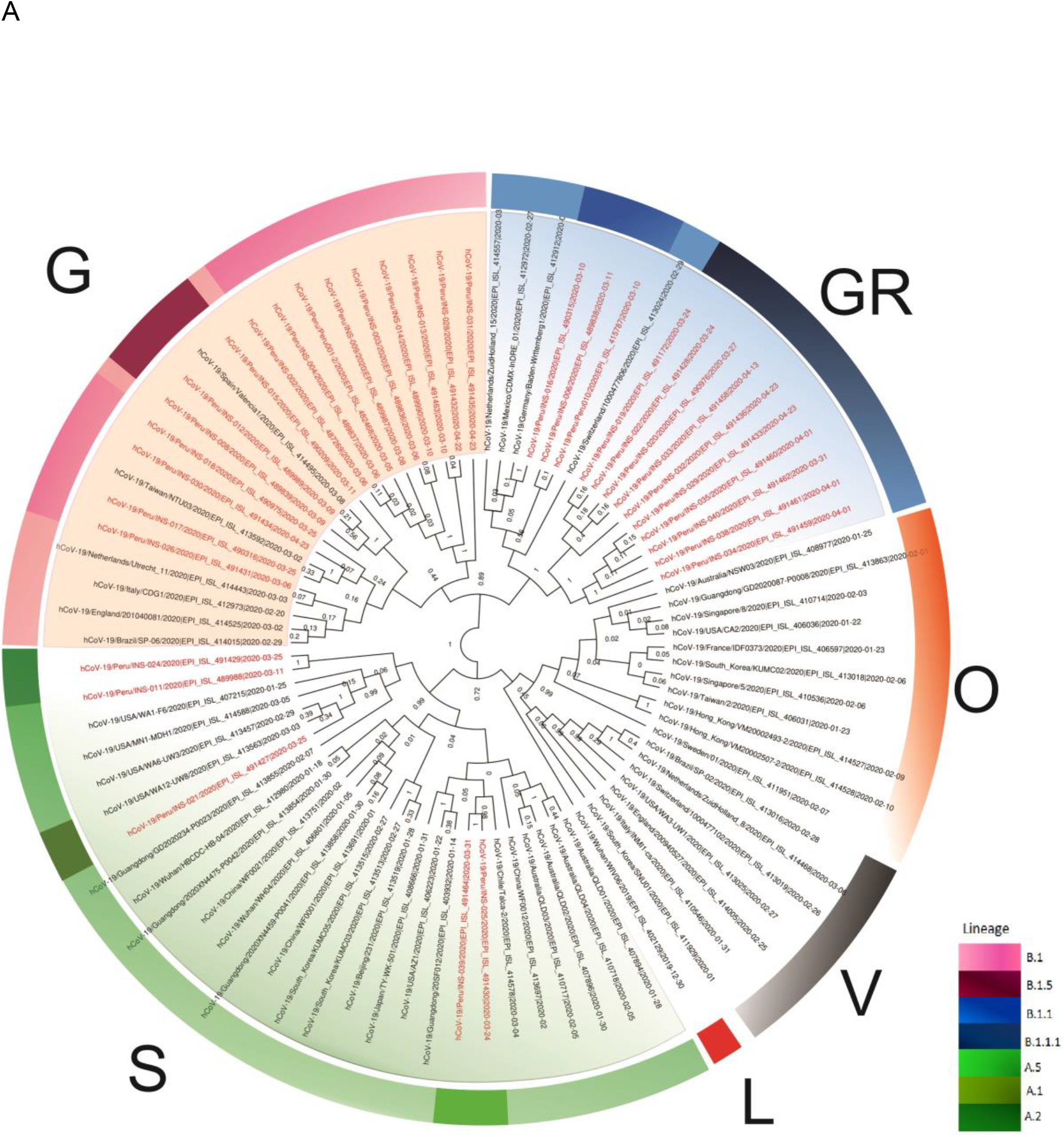
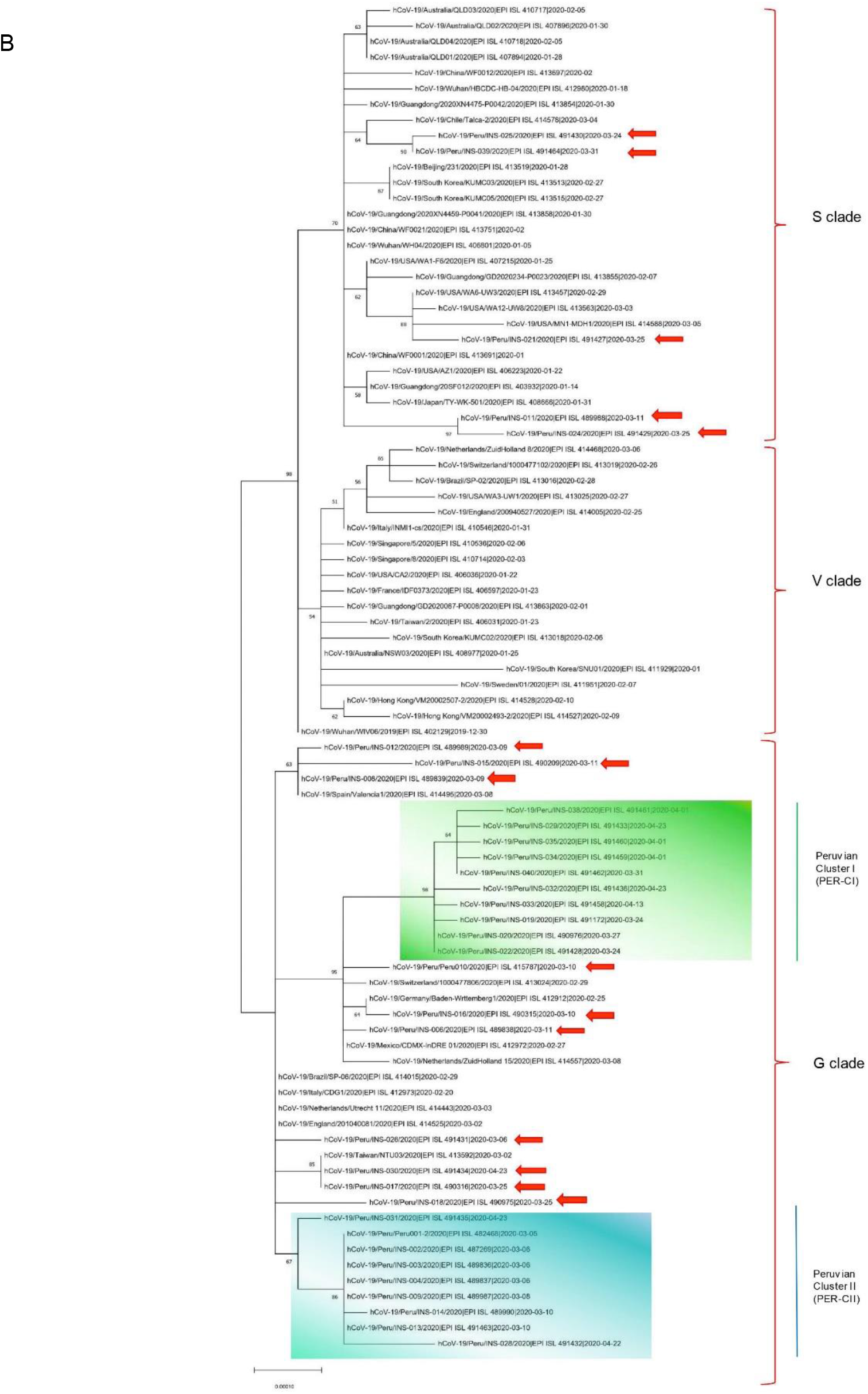
Phylogenetic analysis of SARS-CoV-2 coronavirus isolates. A) Phylogenetic tree built with Beast 2.0. The numbers found in the clades correspond to the posterior probability of each branch. B) Phylogenetic tree built with MEGAX using the Maximum Likelihood algorithm, the numbers in the clades correspond to Bootstrap value. The codes of the Peruvian isolates are in red.

The Peruvian SARS-CoV-2 grouped in clade G presents the characteristic D614G mutation in the Spike (S) protein. The phylogenetic analysis indicates that case zero (LIM-01-2) was closely related to cases reported between 6^th^ and 10^th^ March (LIM-INS-002, LIM-INS-003, LIM-INS-004, LIM-INS-009, LIM-INS-013, and LIM-INS-014) and cases reported in April (LIM-INS-028 and LIM-INS-031) in Lima. We named this cluster as Peruvian Cluster II (PER-CII). Two genomes (LIM-INS-017 and LIM-INS-030) found in Lima were closely related to a genome from Taiwan.

Three genomes from Ica (ICA-INS-012), Lima (LIM-INS-015), and Huánuco (HUC-INS-008), respectively, shown closely related to a case from Spain. The LIM-INS-006, LIM-010, and LIM-INS-016 isolates from Lima were closely related with an isolate from Germany, and with cases reported later (end of March and April) in Lima (LIM-INS-019, LIM-INS-022, LIM-INS-020, LIM-INS-033, LIM-INS-032) and Lambayeque (LAM-INS-034, LAM-INS-035, and LAM-INS-038) and Ica (ICA-INS-040). These samples grouped an almost exclusive cluster (Peruvian Cluster I or PER-CI) shared with European samples. This cluster has the B.1.1.1 or GR lineage, unlike PER-CII that has B.1 o G lineage. We noticed that inside PER-CI cluster exists a subclade formed by samples of Lambayeque and Ica (north and south region of Peru).

The genomes of Peruvian isolates found in clade S have the L84S mutation in the ORF8 region in common. In this clade, we observed that the isolates LIM-INS-025 (Lima) and ICA-INS-039 (Ica) related to a genome reported in Chile. LIM-INS-011 and LIM-INS-024 from Lima related to each other. Simultaneously, LIM-INS-021 from Lima related to genomes reported in the United States.

The MST analysis results showed multiple SARS-CoV-2 introductions to Peru from people who entered the country before the closure of international borders (16^th^ March). We observed the distribution of two main genotypes, the clade S that groups five genomes corresponding to the first notified cases in Peru shortly after the border closure (LIM-INS-021 and ICA-INS039), except for one case (LIM-INS-011) who entered to Peru before that date. The clade G shows more cases (LIM-010 and LIM-01-2, LIM-INS-002, LIM-INS-003, LIM-INS-004, LIM-INS-009, and LIM-INS-013) that entered to Peru before the border closing date (Figure 3).

**Figure 3.**
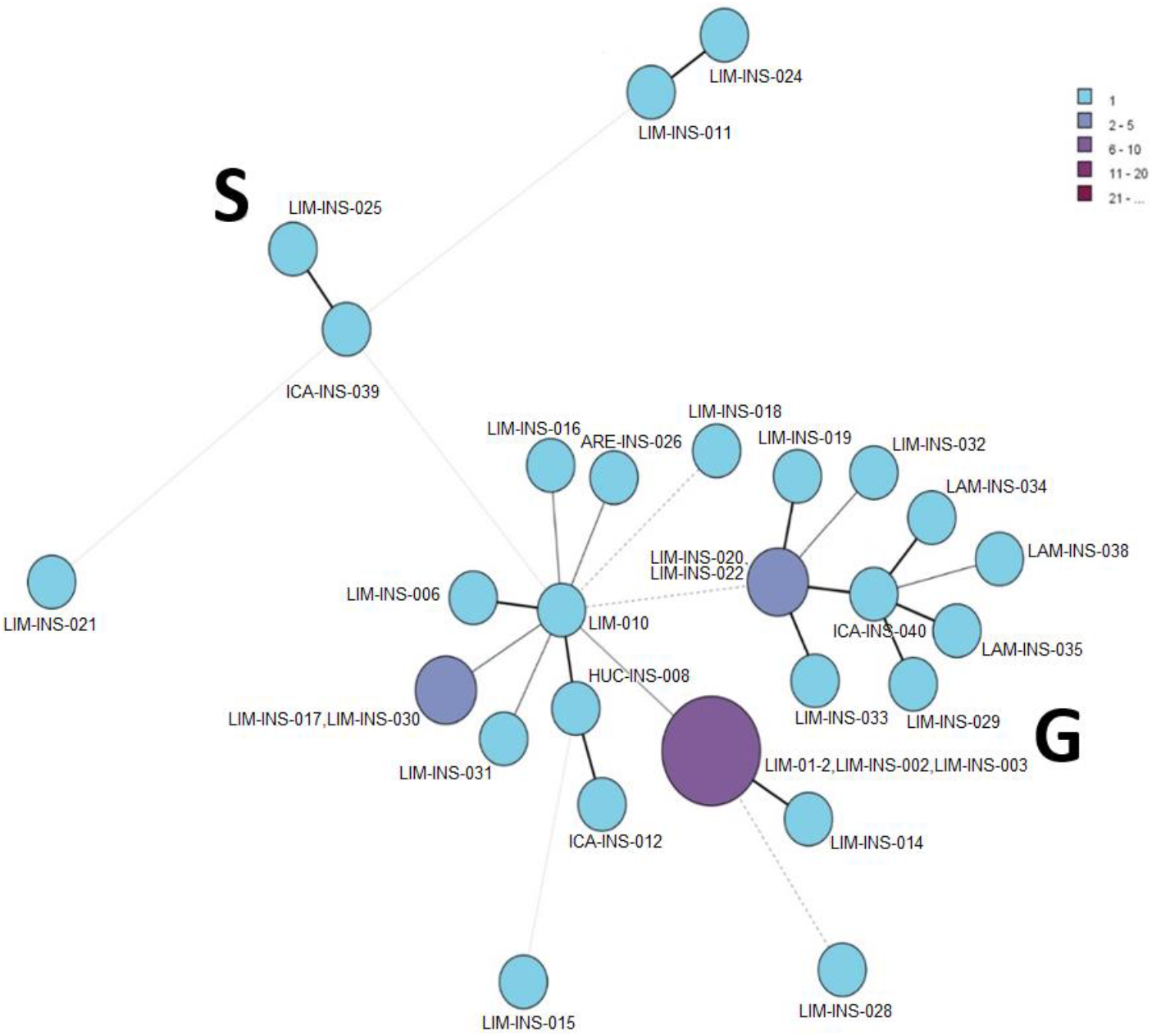
Minimum Spanning Tree (MST) of the first 34 reported genomes. Clades S and G are shown, each showing several introductions occurred before the border closures (Peru11, Peru10, Peru 1, 2, 3, 4, 9, and 13). The other cases show local transmission that began to circulate after the border closures.

**Figure 4.**
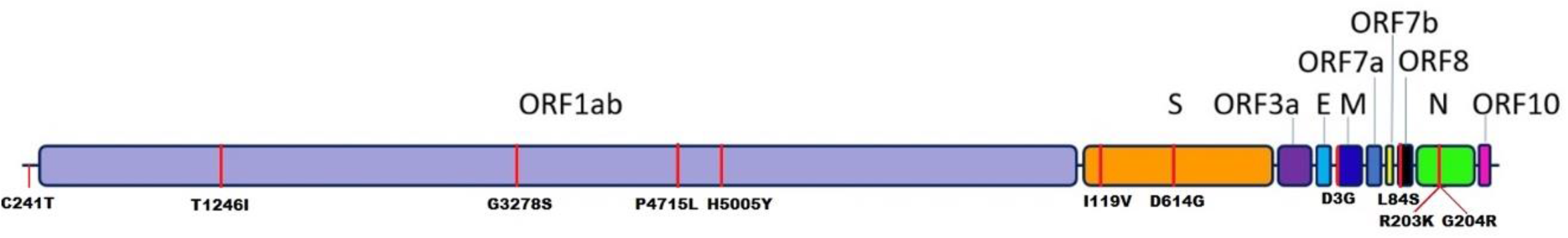
Most frequent variations in the genome of Peruvian SARS-CoV-2 coronavirus isolates. The reference amino acids are on the bar graph, while the variation is below. The scale indicates the size of the SARS-COV2 genome.

The other cases began to circulate after the border closure. These introductions gave rise to other cases, which originated viruses with their unique mutations, generating an autochthonous transmission of the virus.

### Amino acid variations

We found various nucleotide variations in the Peruvian SARS-CoV-2 genomes compared with the reference genome of the Wuhan strain (NC_045512), some of which suffered amino acid changes (Table 3 and S1). We observed the C241T nucleotide variation in the 5’ -UTR region in 30 of 34 genomes.

**Table 3.**
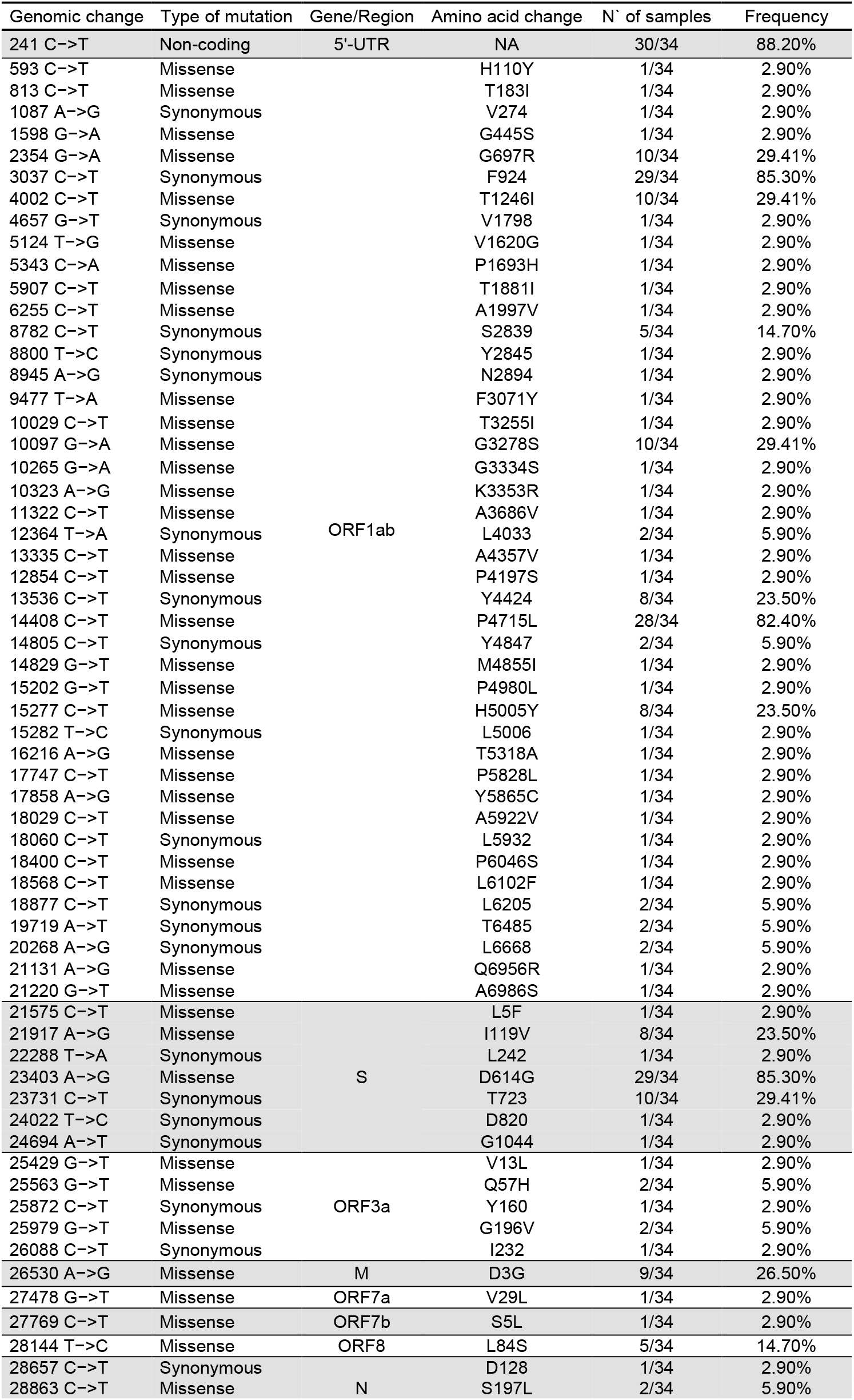
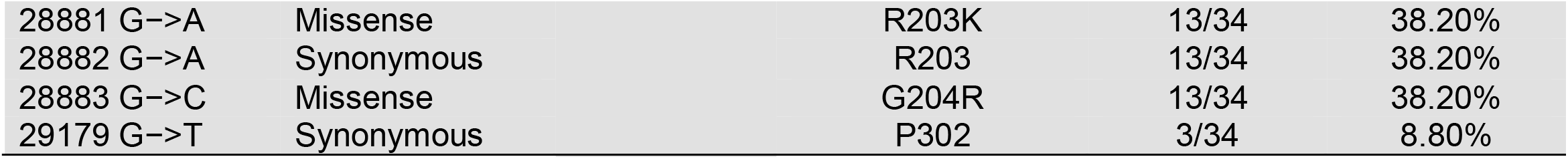
Variations of SARS-Cov2 genomes observed in 34 Peruvian isolates.

Among the most common significant non-synonymous nucleotide variations in ORF1ab are T1246I (10/34), G3278S (10/34), P4715L (29/35), and H5005Y (8/34). We found three amino acid variations in the spike protein: L5F (1/34), I119V (8/34), and D614G (29/34). In the ORF3a region, we found four amino acid variations: V13L (1/34), Q57H (2/34), W149R (1/34), and G196V (2/34). Some genomes present the D3G variation (9/34) in the M protein, a variation found only in samples from the PER-CII cluster. In ORF6 we found the amino acid variation I36T (1/34). Other variations encountered were V29L (1/34) in ORF7a and S5L (1/34) in ORF7b.

In ORF8, we found the L84S variation (5/34), while the protein N presented the following changes, S197L (2/34), R203K (13/34), and G204R (13/34). In this gene, we observed that R203K, G204R, and the synonyms change 28881G>A, are characteristic of PER-CI cluster. We do not find amino acid variations in the gene E and ORF10 (Table 3 and S1). Likewise, we found nucleotide variations in a minority of reads in some positions of 13 sequenced genomes that we do not observe in the final assembled genome (Table 4).

**Table 4:** Nucleotide variations detected in minorities reads. Only variations with coverage greater than 100x and a percentage of reads greater than 10% are recorded.

| Sample | Gene | Nucleotide position | Coverage | Nucleotide variation | % of reads | Amino acid variations |
| --- | --- | --- | --- | --- | --- | --- |
| LIM-INS-013 | S | 22295 | 394 | C -> T | 19.90% | H245Y |
|  | ORF7b | 27792 | 365 | T -> G | 11.60% | F13C |
| LIM-INS-017 | ORF1ab | 9561 | 948 | C -> T | 23.40% | S3099L |
|  | M | 26784 | 2214 | G -> A | 17% | V88I |
| LIM-INS-019 | N | 28738 | 669 | G -> T | 10.90% | A156S |
| LIM-INS-021 | ORF1ab | 12084 | 1334 | C -> T | 31.60% | T3940I |
|  |  | 12085 | 1340 | C -> T | 31.50% | Synonym |
| LIM-INS-022 | ORF1ab | 16433 | 1093 | G -> A | 15.70% | E5457K |
|  |  | 21341 | 253 | A -> T | 28.70% | I7093F |
|  |  | 21342 | 254 | T -> A | 28.30% | I7093N |
|  | S | 22403 | 126 | G -> A | 17.50% | E281K |
|  |  | 22977 | 164 | T -> C | 11% | I472T |
|  |  | 23475 | 266 | C -> T | 17.30% | T638I |
|  | E | 26447 | 735 | C -> T | 12% | S68F |
| LIM-INS-024 | ORF1ab | 18175 | 139 | C -> T | 31.90% | P5971S |
|  |  | 18284 | 158 | A -> T | 34.20% | H6007L |
|  |  | 20047 | 107 | A -> C | 13.10% | I6595L |
|  | ORF10 | 29620 | 1987 | G -> A | 18.50% | M21I |
| LIM-INS-028 | N | 28677 | 508 | C -> T | 29.90% | T135I |
| LIM-INS-029 | ORF1ab | 3139 | 205 | C -> T | 11.20% | S2625F |
| LIM-INS-031 | ORF1ab | 19384 | 295 | G -> A | 12.50% | D6374N |
| LIM-INS-032 | ORF1ab | 14789 | 403 | T -> C | 15.50% | I4842T |
|  |  | 16657 | 695 | C -> T | 21.40% | T5465I |
| LAM-INS-034 | ORF1ab | 596 | 569 | G -> T | 10.90% | V111L |

## DISCUSSION

At the end of December 2019, discovering a new virus that could cause respiratory disease in humans was announced in Wuhan-China. At that time, we all thought that the virus would take a long time for it to spread to other continents; however, after three months, the first case was reported in Peru, and from that time to the present, we continue with more positive cases per day being difficult to control this pandemic.

In this context, it is necessary to research to know the genomic diversity of SARS Cov2 in different regions of the country and share this data with scientists interested in developing diagnostic methods and possible vaccines.

In this study, we report the main clades and genomic lineages of 34 genomes of SARS-CoV-2 virus from different regions, at the beginning of the pandemics in Peru. A previous inspection of SARS-CoV-2 genomic sequences from South America available on GISAID revealed that the lineage B.1 is the most prevalent (82%) SARS-CoV-2 variant circulating in South America (23). Our results show that the B.1 lineage and its sub-lineages were also the most prevalent in Peruvian analyzed areas (n=29, 85.3%).

Phylogenetic analysis indicates that the SARS-CoV-2 virus in Peru belongs to the genetic clades G and S. Both clades entered Peru almost simultaneously according to the date of sample collection. The first confirmed case in Peru in early march was a man infected during their visit to European countries; this hypothesis is confirmed by both phylogenetic analysis and the patient’s epidemiological data. Later, in late March and early April, we observed the appearance of new cases genetically related to the first reported cases. This observation is compatible with a local transmission since the Peruvian government decreed the closure of borders and a state of emergency in mid of march. This same pattern of events was reported in other countries in America (11, 23, 24).

The B1.1.1 lineage found in Peru would be responsible for most local transmission cases and derived in two Peruvian sublineages (PER-CI and PER-CII). These sublineages present a haplotype of two non-synonymous mutations in PER-CI (R203K and G204R), and one synonymous mutation (28881G>A) in Nucleocapsid protein; which is similar to lineages B.1.1.EU/BR and B.1.1.BR found in Brazil (23); and one mutation for PER-CII (D3G in M gene) (Table S1).

We observed several mutations at different positions in the genome of circulating SARS-COV-2. It is worth highlighting the presence of the C241T (30/35) mutation in the 5’ UTR region important for transcription and viral packaging (12, 13). The P4715L mutation (14408 C>T) in the viral polyprotein ORF1ab located in the RNA-dependent RNA polymerase region (position 323 of RdRP), which could be related to the increase in the mutation rate (14). This polymerase dynamics simulation indicates that this P4715L is a more stable RdRP variant (15). The spike protein’s D614G mutation is related to the increase of the virus’s infectivity and is present in 29 of the 35 genomes reported in Peru (16). Likewise, it is remarkable that the I119V mutation (present in 8 from the 34 genomes) is present in the N-terminal domain (NTD) of the S1 region of the spike, and we detected no variations in the RBD region. Meanwhile, the R203K (13/34) and G204R (13/34) variants of the N gene could contribute to the reduction of the nucleocapsid’s conformational entropy (17). ORF3a is a carrier for K+ and Ca2+ ions (18) and is necessary for *in vitro* and *in vivo* replication (19). The Q57H mutation detected in ORF3a could generate a constriction in the formed pore (20).

Besides, we found some viral subpopulations or quasi-species, defined by the presence of mutations in a percentage of detected reads, consistent with that previously reported for SARS-COV-2 (21) and influenza virus (22). These mutations that appear over time are likely due to the virus’s adaptation to the new human host. Likely, these mutations will not fix or generate a new variant unless these variations could improve the virus’s adaptation to its host or increase the virus’s virulence.

An important limitation of our study is the uneven spatial and temporal sampling scheme, most SARS-CoV-2 sequences recovered in the present study were from Lima and a few samples of north and south regions of Peru. These sampling does not represent the viral diversity in other regions of Peru. More accurate reconstructions of the origin and regional spread of the clade B.1.1.1 will require a denser sampling from Peru and neighboring South American countries, particularly during the very early phase of the epidemic. It is crucial to analyze more cases by whole genome sequencing of the SARS-CoV-2 virus from different regions of the country, to better understand the spread and evolution of this new coronavirus in Perú. It is also important to integrate the information from the laboratory diagnosis and the genomic analysis of the virus with the clinical and epidemiological data to depict the clinical epidemiology and infection mechanisms of this coronavirus in Peru.

## MATERIALS AND METHODS

Nasal and pharyngeal swabs summited to the National Reference Respiratory Viruses Laboratory of the Peruvian National Institute of Health for diagnostic confirmation were stored at −80 °C. We selected samples from the first cases reported in Peru and from different regions of the country. The first 34 samples, from early March to early April, were chosen following the criteria that they had an adequate viral load (Ct <30) after analyzing the molecular diagnosis results by real-time RT-PCR.

Viral RNA was purified from 140 μL of viral transport medium, using the QIAamp Viral RNA Mini Kit (Qiagen, Germany) following the manufacturer’s instructions. The purified RNA samples were processed by reverse transcription and PCR to amplify overlapping fragments, using primers designed (Table 1) with the PrimerQuest Tool software (www.idtdna.com) using the SARS-COV-2 genome from a Wuhan isolate (Genbank accession number NC_045512. 2) and a pool of genomes reported so far. The amplification process was done using SuperScript ™ IV One-Step RT-PCR System kit (Invitrogen) as follows: 12.5 μL of 2X amplification buffer, 1.25 μL of forward primer at 10 μM, 1.25 μL of reverse primer at 10 μM, 4.5 μL of PCR water, 0.5 μL SuperScript IV / High Fidelity polymerase and 5 μL of purified RNA. The amplification program: 50 °C for 45 minutes, 98 °C for 2 minutes followed by 45 cycles of 98 °C for 10 seconds, 54 °C for 30 seconds and 68 °C for 5 minutes, finally an extension of 68 °C for 10 minutes. The amplification products purified with the PureLink ™ PCR Purification kit (Invitrogen).

The amplified fragments (2 ng) were processed using the Nextera XT DNA Library Prep kit and indexes adapters (Illumina) following the procedure recommended by the manufacturer. The library was sequenced using a Nextera XT DNA Library Prep Kit, a MiSeq Reagent Kit v2 cartridge (500-cycles), and the MiSeq sequencer (Illumina). The fastq files generated were cleaned using Groomer and Trimmomatic (https://usegalaxy.org/). The reads mapped to the Wuhan-Hu-1 reference genome available at the National Center for Biotechnology Information (https://www.ncbi.nlm.nih.gov/nuccore/?term=Severe+acute+respiratory+syndrome+coronavirus+2+AND+Wuhan) using BWA-ME (https://usegalaxy.org). Finally, the genomes were assembled using SPAdes and compared to the reference genome using CONTIGuator (http://contiguator.sourceforge.net/). Nucleotide and amino acid variations were detected using snpEff (http://snpeff.sourceforge.net/SnpEff_manual.html). Likewise, a manual edition of the genomes was done using the Geneious Prime software (https://www.geneious.com/).

The phylogenetic analysis of the genomes was done using the Bayesian Markov Chain Monte Carlo method (MCMC) using BEAST 2 (9). The Yules demographic model and the Log-Normal Relaxed clock were selected, running 10 MCMC chains of 10 million. Convergence was evaluated with Tracer (https://www.beast2.org/tracer-2/), obtaining an ESS higher than 200. The Maximum Clade Credibility (MCC) tree is generated after a 10% burning using TreeAnnotator. The MCC tree was visualized and edited using FigTree (http://tree.bio.ed.ac.uk/software/figtree/). Additionally, to detect local transmission of the virus we performed Maximum Likelihood analysis with 1000 bootstraps of confidence using MEGA X (https://www.megasoftware.net). For the identification of the SARS-CoV-2 genotypes or genetic lineages, we analyzed the 34 genomes using the recently described Pangolin 2.0 web application (http://cov-lineages.org) (10).

A minimum spanning tree (MST) analysis using BioNumerics version 7.6 (Applied Maths NV, Sint-Maartens-Latem, Belgium) drawn evolutionary relationships between samples. MST connects each genome based on the degree of changes required to go from one allele to another. The MST structure is represented by branches (continuous vs. dashed and dotted lines) and circles representing each pattern. The length of the branches represents the distance between patterns; while the complexity of the lines (continuous, gray dashed and gray dotted) denotes the number of SNP changes between two patterns: solid lines, 1 or 2 or 3 changes (thicker ones indicate a single change, while the thinner ones indicate 2 or 3 changes); gray dashed lines represent four changes; and gray dotted lines represent five or more changes. The size of the circle is proportional to the total number of genomes.

## Acknowledgments

The authors thank all the health personnel who heroically and hard face the effects of the COVID-19 disease in Peru. Especially the staff of the National Institute of Health and the COVID-19 diagnostic team of INS Peru. In addition, we thank Sr. Harrison Montejo for his contribution to the achievement of the objectives of this project.

This research work has been funded by the National Institute of Health of Peru.

## Contributions

CPR, KVCh, MGP, HBC, PLP, JBT, MHN, DGN, NRS, OCR contributed with Study design, Data collection, Data analysis, and Writing. KVCh, CPR, MGP and DGN contributed with samples processing and sequencing. All authors agree with the final version of the article.

## Supporting information

**Table S1:**
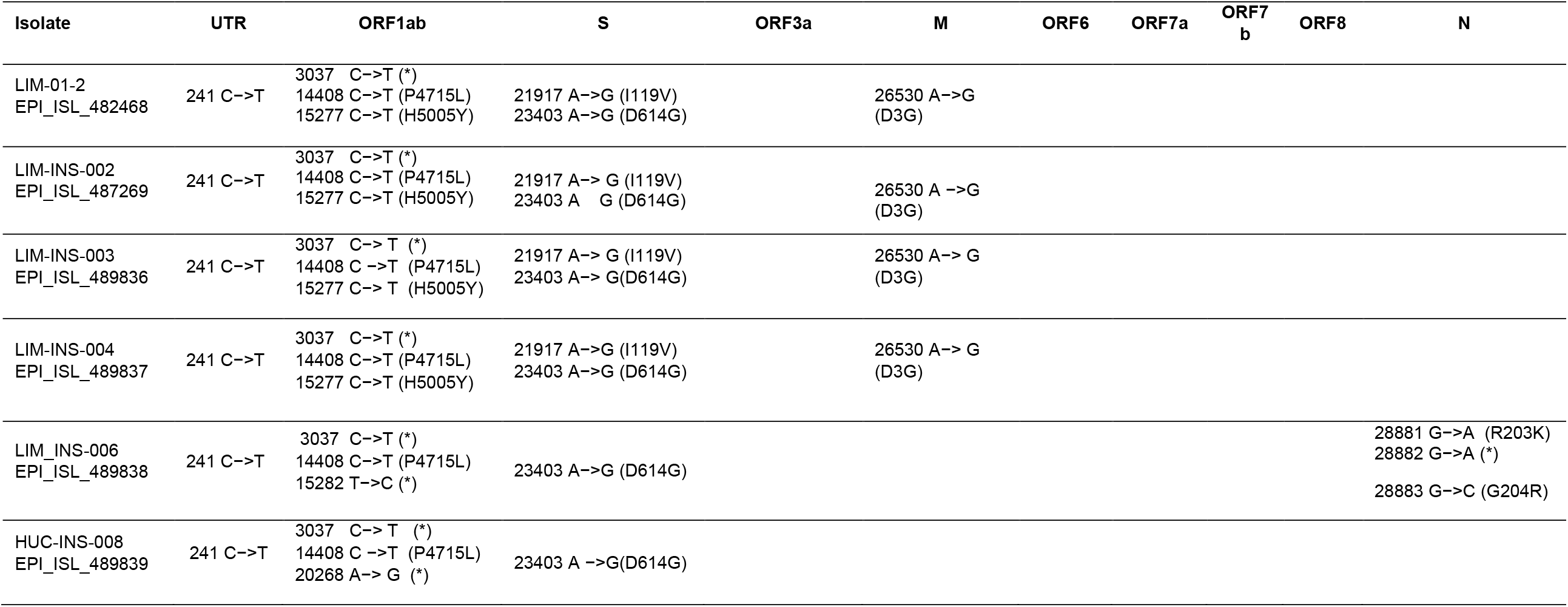
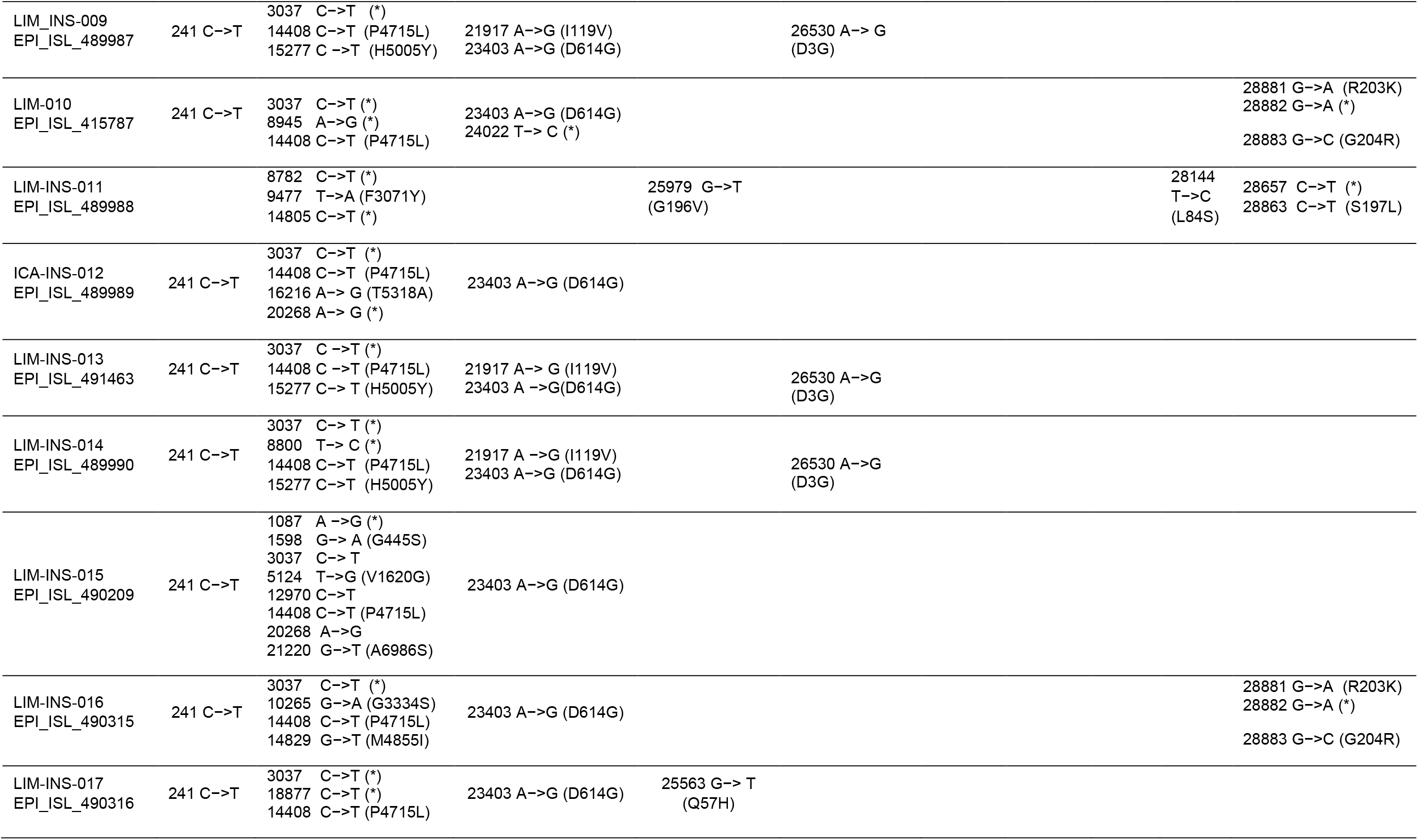
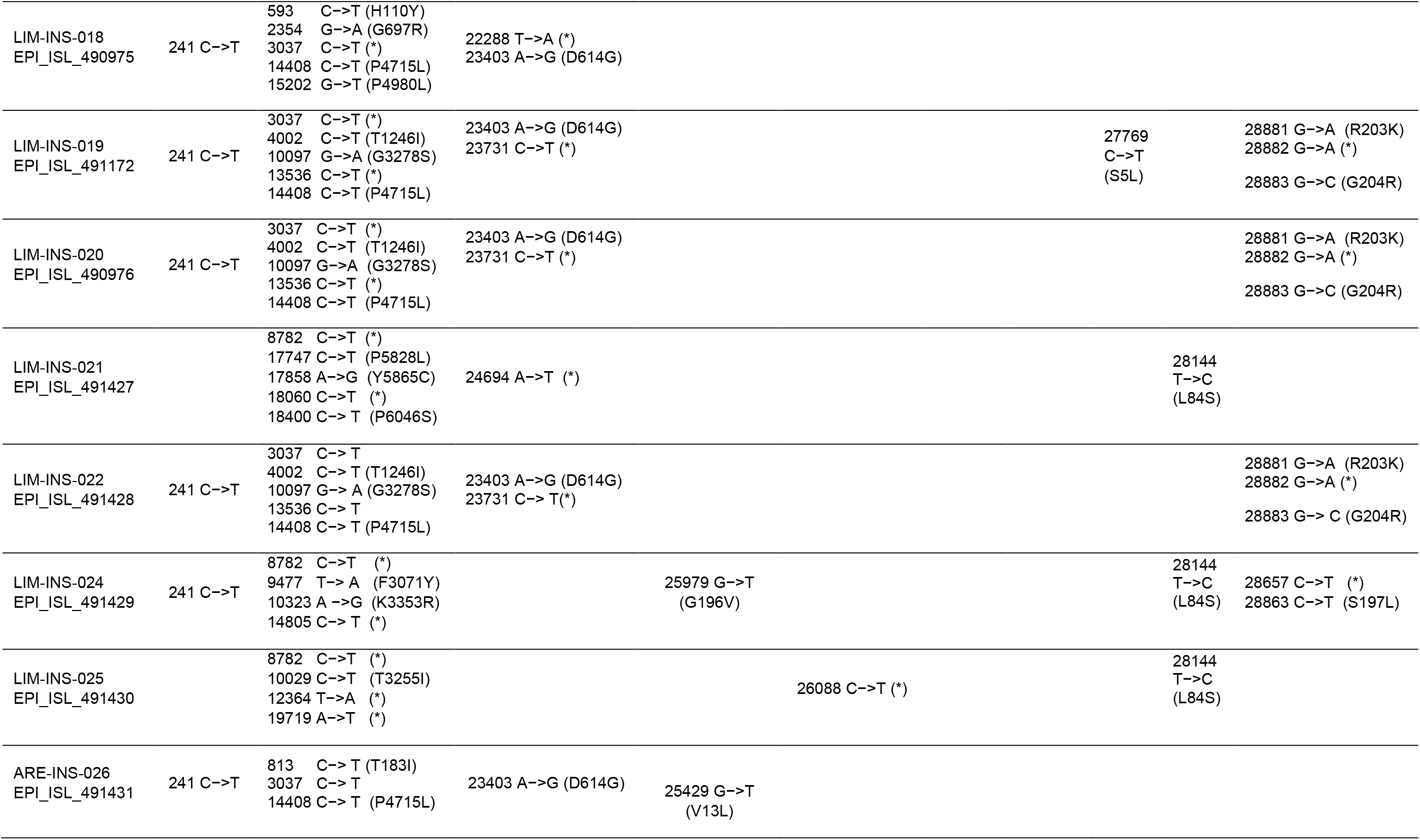
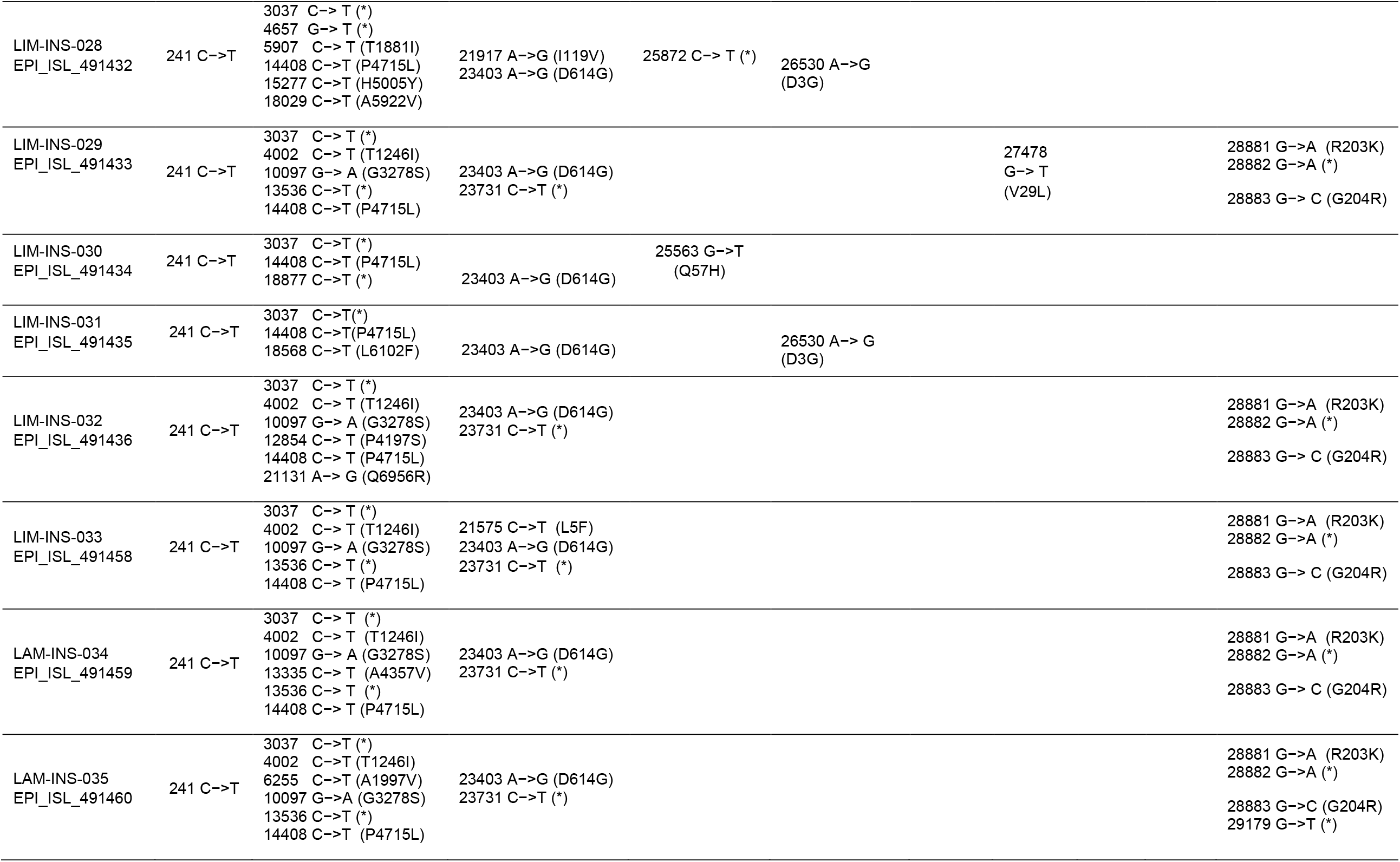
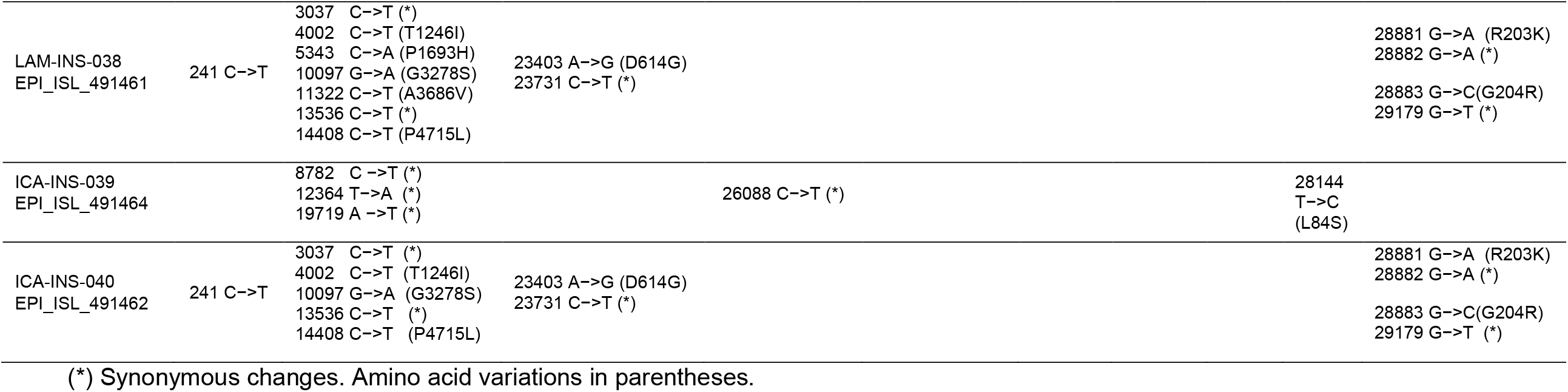
Nucleotide and amino acid variations in the genomes of sequenced Peruvian isolates.

